## Supplementary material for "Evolution of bacterial persistence to antibiotics during a 50,000-generation experiment in an antibiotic-free environment": Figure S1- Effect of antibiotics on growth rates

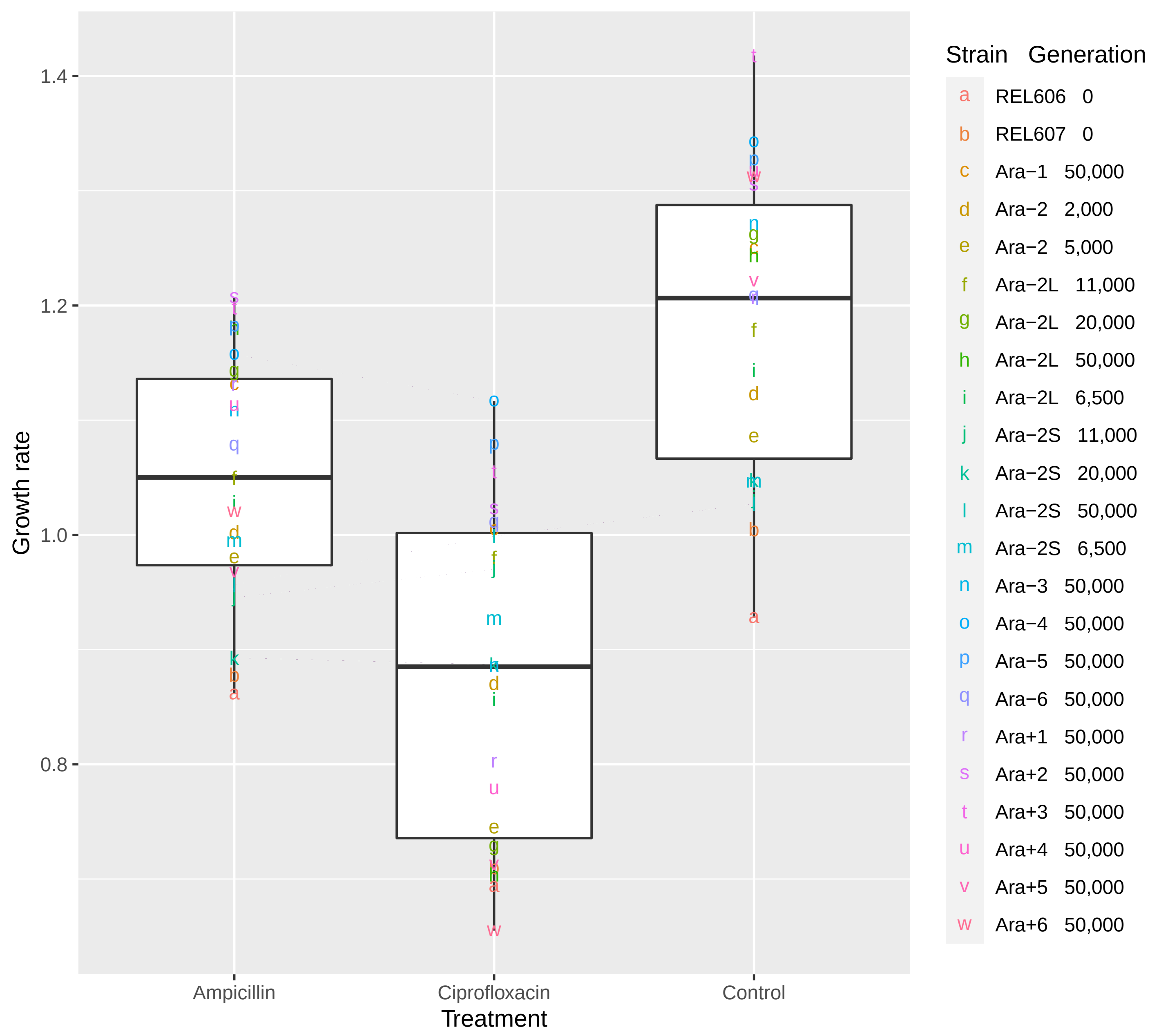


Figure S1: Effect of antibiotics on growth rates.
Each combination of letter and colour corresponds to one clone.
Lines link each individual clone in each treatment.
In the boxplots: the central line give the median, the extremities the first and third quartiles, and the vertical bars the minimal and maximal values.
