## Supplementary material for "Evolution of bacterial persistence to antibiotics during a 50,000-generation experiment in an antibiotic-free environment": Table S1- Tukey Comparisons - Only Significant

| **Ara** | -2L | -2S | -3 | -4 | -5 | -6 | +2 | +3 | +4 | +5 | +6 |
| --- | --- | --- | --- | --- | --- | --- | --- | --- | --- | --- | --- |
| -2L |  | ln(FC) = 5.81  p < 0.001 *** |  | ln(FC) = 5.58  p = 0.016 * |  |  | ln(FC) = 5.51  p = 0.013 * |  |  |  |  |
| -2S |  |  |  |  |  |  |  |  |  |  |  |
| -3 |  |  |  |  |  |  |  |  |  |  |  |
| -4 |  |  |  |  |  |  |  |  |  |  |  |
| -5 |  |  |  |  |  |  |  |  |  |  |  |
| -6 |  |  |  |  |  |  |  |  |  |  |  |
| +2 |  |  |  |  |  |  |  |  |  |  |  |
| +3 |  |  |  |  |  |  |  |  |  |  |  |
| +4 |  | ln(FC) = 9.57  p < 0.001 *** | ln(FC) = 6.76 ; p < 0.001 *** | ln(FC) = 9.34  p < 0.001 *** | ln(FC) = 6.23  p = 0.009 ** | ln(FC) = 7.83  p < 0.001 *** | ln(FC) = 9.27  p < 0.001 *** | ln(FC) = 6.39  p = 0.002 ** |  |  |  |
| +5 |  | ln(FC) = 6.6  p = 0.001 ** |  | ln(FC) = 6.37  p = 0.036 * |  |  | ln(FC) = 6.29  p = 0.034 * |  |  |  |  |
| +6 |  | ln(FC) = 5.71  p < 0.001 *** |  | ln(FC) = 5.48  p = 0.03 * |  |  | ln(FC) = 5.4  p = 0.028 * |  |  |  |  |

Table S1: Pairwise comparisons of the abundance of ciprofloxacin persister cells in the evolved clones sampled from each of the 12 LTEE populations at 50,000 generations.

These comparisons are based on the coefficients of the model in Table 1. The ln(FC) values give the logarithm of the fold change of the abundance of persister cells. This fold change is defined as the abundance in the clone in the row over the abundance in the clone in the column. In non-empty cells, persistence is significantly higher in the clone in the row than in the clone in the column. The significance of this fold change was assessed with the R package multcomp. For readability, the clones sampled from populations Ara‒1 and Ara+1 were removed because they do not significantly differ from any other clones, and only significant results are shown. For the full results, see tables S2 and S3. Significance codes: 0 ‘***’ 0.001 ‘**’ 0.01 ‘*’ 0.05 ‘.’ 0.1 ‘ ’ 1
