## Supplementary material 1- Heuristic detection of the exponential growth phase for "Evolution of bacterial persistence to antibiotics during a 50,000-generation experiment in an antibiotic-free environment": 0_Description of the heuristic selection of the exponential growth phase.docx

We detected the exponential growth phase in two steps that were implemented by the R function “*SimpleExponentialGrowthFit*” that we developed (file *SimpleExponentialGrowthFit.R)*. The rationale is similar to the approach implemented by the R function ‘*fit_easylinear*’ (details and comparison below).

In the **first** step, we identified the part of the growth curve with the highest exponential link between OD_600_ and time by using a sliding window approach. While ‘*fit_easylinear*’ implicitly assumes that the background was removed by the user, our function estimates the background separately for each growth curve. This is important since the background vary a lot within the same plate, as shown in file ‘*heterogeneity between initial OD.pdf*’. Then to consider the entire exponential growth phase, the **second** step aimed at extending the span of the window with the highest exponential link between OD_600_ and time that was detected during the first step.

We detected and fitted exponential growth for each growth curve as follow.

**Step 1: finding the window with the highest exponential link between OD_600_ and time.**

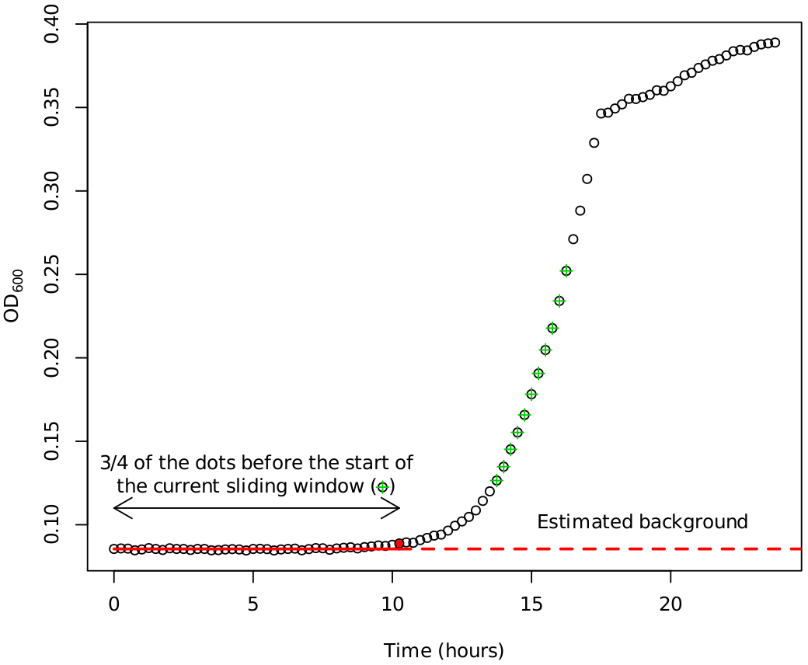
For each sliding window of 10 time points (2.5h) we fitted and scored the linear model
${{log}_{2}\left( OD-Bg \right)}_{i}=GrowthRate\times{time}_{i}+{OD}_{init}+\varepsilon_{i}$ where:

- $Bg$ is the background OD that is specific to the sliding window. It is estimated as the median OD in the first ¾ of the dots before the start of the current sliding window (e.g. in Figure S2).
- ${time}_{i}$ is the explanatory variable.
- $GrowthRate$is the slope associated with this variable. Indeed, the time needed for $OD-Bg$ to be multiplied by two is$1/GrowthRate$ . To see this, define ${OD}_{b}$such as${OD}_{b}-Bg=2\times{OD}_{b}-Bg \Leftrightarrow{log}_{2}\left( {OD}_{a}-Bg \right)={log}_{2}\left( 2 \right)+{log}_{2}\left( {OD}_{b}-Bg \right)=1+{log}_{2}\left( {OD}_{b}-Bg \right)$. This equation can be rewritten with the right side of the model: $GrowthRate\times{time}_{b}+RSG=1+GrowthRate\times{time}_{a}+RSG\Leftrightarrow GrowthRate\times\left( {time}_{b}-{time}_{a} \right)=1\Leftrightarrow{time}_{b}-{time}_{a}=1/GrowthRate$.
- ${OD}_{init}$ is the intercept that corresponds to the initial OD. Indeed, at${time}_{1}=0$, we have${OD}_{0}-Bg=2^{{OD}_{init}}$. In the absence of lag phase, this value would correspond to $a\times N_{0}$ where $N_{0}$ is the initial number of cells, and $a$ describes the relationship between $OD$ and $N$: $OD-Bg=a\times N$. However, since a lag phase is present in most cases, $2^{{OD}_{init}}$ is actually positively associated with $N_{0}$ and negatively to the lag phase.

Figure S2: Illustration of the detection of the exponential phase in the growth curve.

We scored the model of each sliding window with the formula:

$$GrowthRate\times ln(standard deviation of the OD in the conscidered sliding window+1)\times R^{2}$$

and selected the sliding window with the highest score.

In this formula, the $ln$ of the OD's standard deviation avoids that the window is selected earlier than the exponential growth, i.e., during the lag phase, when growth revealed to be noisy.

**Step 2: extending the span of the window.**

We extended the window span to ensure it includes the entire exponential phase. We chose the span of the window that maximized $-\bar{log\left( \left| \hat{OD}-OD \right| \right)}\times R^{2}$ where $\hat{OD}$ is the predicted OD by extrapolating the model adjusted to the selected window to the entire growth curve. The use of the log of the absolute value of the error allowed to set a low weight to large errors that corresponded to the growth period outside the exponential growth phase.
