## Supplementary material 1- Heuristic detection of the exponential growth phase for "Evolution of bacterial persistence to antibiotics during a 50,000-generation experiment in an antibiotic-free environment": SimpleExponentialGrowthFit.R

```
SimpleExponentialGrowthFit =function(DO, Time
                               ,ExploredTime=NULL
                               ,NdotsFirtsFit=ifelse(is.null(ExploredTime),5,length(Time[(Time >= ExploredTime[1]) & (Time <= ExploredTime[2])]))
                               ,Plot=TRUE,Log=FALSE,Main=NULL){
  # 
  # library(lmtest)
  #
  Time_ = Time
  DO_ = DO
  if(!is.null(ExploredTime)){
    if(max(Time) < ExploredTime[2]){stop("max(Time) < ExploredTime[2]")}
    if(min(Time) > ExploredTime[1]){stop("min(Time) > ExploredTime[1]")}
    #
    DO   = DO[  (Time_ >= ExploredTime[1]) & (Time_ <= ExploredTime[2])]
    Time = Time[(Time_ >= ExploredTime[1]) & (Time_ <= ExploredTime[2])]
  }else{ExploredTime = range(Time_)}
  #
  # RM = seq(0, min(DO)-min(DO)/100 ,length.out = 100)
  # RM = seq(0, min(DO)-min(DO)/100 ,length.out = 5)
  # RM = seq(0, (max(DO)-min(DO))/5+min(DO) ,length.out = 5)
  # for(tuneRMiteration in 1:10){
    END = NdotsFirtsFit:length(Time)
    SlopFirstFits = 
      # sapply(RM,function(rm){
      sapply(END,function(end){  # print(c('rm' = rm, 'end' = end))
        W = (end-NdotsFirtsFit+1):end
        rm = median(DO[1:(round(min(W)/4*3))])
        DOlog2 = suppressWarnings(log2(DO-rm))
        DOlog2[is.infinite(DOlog2)] = NA
        while(sum(!is.na(DOlog2[W])) < NdotsFirtsFit*0.666){
          rm = rm*0.99
          DOlog2 = suppressWarnings(log2(DO-rm))
          DOlog2[is.infinite(DOlog2)] = NA
        }
        # if(sum(!is.na(DOlog2[W])) < NdotsFirtsFit*0.666  ){-Inf}else{
          Range = range(W)
          M = lm(DOlog2[W] ~ Time[W])

          coef(M)[2]*
          log(sd(DO[W])+1) *
          summary(M)$r.squared
          # }
        # })
      # plot(DOlog2 ~ Time)
    })
    end = END[ which.max(SlopFirstFits)]
    W = (end-NdotsFirtsFit+1):end
    rm = median(DO[1:(round(min(W)/4*3))])
    DOlog2 = suppressWarnings(log2(DO-rm))
    DOlog2[is.infinite(DOlog2)] = NA
    while(sum(!is.na(DOlog2[W])) < NdotsFirtsFit*0.666){
      rm = rm*0.99
      DOlog2 = suppressWarnings(log2(DO-rm))
      DOlog2[is.infinite(DOlog2)] = NA
    }

    # RM = seq(min(RM[order(SlopFirstFits,decreasing = TRUE)[1:3]]), max(RM[order(SlopFirstFits,decreasing = TRUE)[1:3]]) ,length.out = 7)
    # print(rm)
  # }
  DOlog2  = suppressWarnings(log2(DO -rm)) ; DOlog2[ is.infinite(DOlog2 )] = NA
  DOlog2_ = suppressWarnings(log2(DO_-rm)) ; DOlog2_[is.infinite(DOlog2_)] = NA
  # DOlog2 = suppressWarnings(log2(DO-rm)) ; plot(DO~Time) ; points(DO[W]~Time[W],pch=4) ; M=lm(DOlog2[W] ~ Time[W]) ; points(I(2^(coef(M)[1]+coef(M)[2]*Time) + rm) ~ Time, type='l',col=3)
  
  TimeMax = which.min(diff(diff(DO)))
  if(TimeMax < max(W)){TimeMax = max(W)}
  
  if(sum(!is.na(DOlog2[W])) > NdotsFirtsFit*0.666){
    Range = range(W)
    M = lm(DOlog2[W] ~ Time[W])
    Pred = coef(M)[1] + coef(M)[2] * Time
    Err = abs((DO-rm)  -  2^Pred) / diff(range(DO))
    w = FALSE
    div = 4
    while(!sum(w)>5){ div=div/2
      w = Err<diff(range(DO))/div
    }
    Score = -mean(log(Err[w]+1))*
      # coef(M)[2]*
      summary(M)$r.squared #-log(summary(M)$coefficients[2,4])*MaximizeTheNoOfDots  + log(bgtest(I(Pred - DOlog2[Wpred]) ~ Wpred)$p.value)
    #
    improved = TRUE
    while(improved){
      W_ = vector(mode = 'list',length = 9)
      # Slop_ = c()
      Score_ = c()
      i = 0
      Range = range(W)
      # print(Range)
      for(low in 0:3){
        for(up in 0:3){
          if(!(up == 0 & low == 0)){i=i+1
          W_[[i]] = c((Range[1]-low):(Range[2]+up))
          W_[[i]] = W_[[i]][W_[[i]]>0 & W_[[i]]<TimeMax]
          if(min(W) > 0 & max(W)<=TimeMax){
            W = W_[[i]]
            M = lm(DOlog2[W] ~ Time[W])
            Pred = coef(M)[1] + coef(M)[2] * Time
            Err = abs((DO-rm)  -  2^Pred) / diff(range(DO))
            Score_[i] = -mean(log(Err[Err<diff(range(DO))/div]+1)) *
              # coef(M)[2]*
              summary(M)$r.squared #-log(summary(M)$coefficients[2,4])*MaximizeTheNoOfDots  + log(bgtest(I(Pred - DOlog2[Wpred]) ~ Wpred)$p.value)
            if(is.na(Score_[i])){Score_[i] = -Inf}
          }else{
            Score_[i] = -Inf
          }}}}
      # Ndots_ = sapply(W_, length)
      # Score_ = (Ndots_/length(Time))*MaximizeTheNoOfDots + Slop_
      improved = any(Score_ > Score)
      if(improved){
        Score = max(Score_)
        W = W_[[which.max(Score_)]]
      }}
  }else{
    R2 = 0
    GrowthRate=NA
    DOmax = NA
  }
    M = lm(DOlog2[W] ~ Time[W])
    R2 = suppressWarnings(summary(M)$r.squared)
    if(is.na(R2)){R2=0}
    GrowthRate = coef(M)[2]
    DOmax = max(DO)
    if(R2>0.5){
      Col = rep("black",length(Time_)) ; Col[which((Time_ >= ExploredTime[1]) & (Time_ <= ExploredTime[2]))[W]] = 'red'
    }else{
      Col = rep("grey40",length(Time_))
    }
    select = which(Col=='red')
    Pch = rep(1,length(Time_)) ; Pch[select] = 19
    Pred = coef(M)[1] + coef(M)[2] * seq(min(Time_),max(Time_),length.out = 100)
    # Exponential = bgtest(M)$p.value
  # LagPhase
  if(R2>0.7){
    X = seq(-2*Time_[max(select)],Time_[max(select)],length.out = 500)
    w = 1:max(c(1,round(select[1]*2/3)))
    if(length(w)>1){  BaseDOlog2 = log2(  median(DO_[w])  -rm)
                      LagPhase = (BaseDOlog2-coef(M)[1][[1]])/coef(M)[2][[1]]
              }else{  BaseDOlog2 = LagPhase = NA  }
  }else{
    BaseDOlog2 = LagPhase = NA
  }
  if(is.infinite(BaseDOlog2)){BaseDOlog2 = NA}
  if(is.infinite(LagPhase)){LagPhase = NA}
  # 
  if(Plot){
    plot(list(DO_,DOlog2_)[[Log+1]] ~ Time_, ylab = c("DO","DOlog2")[Log+1],xlab = "Time",col=Col,pch=Pch,main=Main)
    if(Log){
      abline(M,col=3)
      points(LagPhase,BaseDOlog2,pch=4,cex=1.5,lwd=2,col=4)
    }else{
      points(seq(min(Time_),max(Time_),length.out = 100),  2^Pred+rm,  type='l',col=3)
      points(LagPhase,2^BaseDOlog2+rm,pch=4,cex=1.5,lwd=2,col=4)
    }
    mtext(paste0(" R²="         ,signif(R2,digits = 2)
                 # ," Pvalue(H0:Exponential)="  ,signif(Exponential   ,digits = 2)
                 ,"\nDOmax="     ,signif(DOmax               ,digits = 2)
                 ,"; GrowthRate=",signif(GrowthRate          ,digits = 2)
    ),side = 3,line = 0.095,cex=0.5)
  }
  return(list("GrowthRate"=GrowthRate
              , "DoublingTime"=1/GrowthRate
              ,"log2_a.N0"=coef(M)[1]
              #' log2(DO) = ax+b
              #' DO = 2^(ax+b) = 2^(ax) * 2^b
              #' We define x0 and x2 and their corresponding values DO0 and DO2 such as DO2 = DO0*2. Therefore x2-x0 is the doubling time
              #          DO2/DO0  =      2      = (2^(a*x2) * 2^b) / (2^(a*x0) * 2^b) = (2^(a*x2)) / (2^(a*x0)) = 2^(a*x2-a*x0) = 2^(a*(x2-x0))
              # <=> log2(DO2/DO0) = log2(2) = 1 = a*(x2-x0)
              # <=> (x2-x0) = 1/a The doubling time
              , "DOmax"=DOmax, 'LagPhase' = c('BaseDOlog2'=BaseDOlog2, 'LagPhase'=LagPhase)
              # , 'Exponential'=Exponential
              , 'R2'=R2, "Col"=Col, "M"=list(M), 'Pch'=Pch, 'Pred'=Pred, 'rm'=rm, 'DO'=DO_, 'DOlog2'=DOlog2_,'Time'=Time_, 'Ndots' = length(W)))
}
```
