## Supplementary figures and images for "Evolution of bacterial persistence to antibiotics during a 50,000-generation experiment in an antibiotic-free environment"

### Figure explaining detection growth curve.pdf

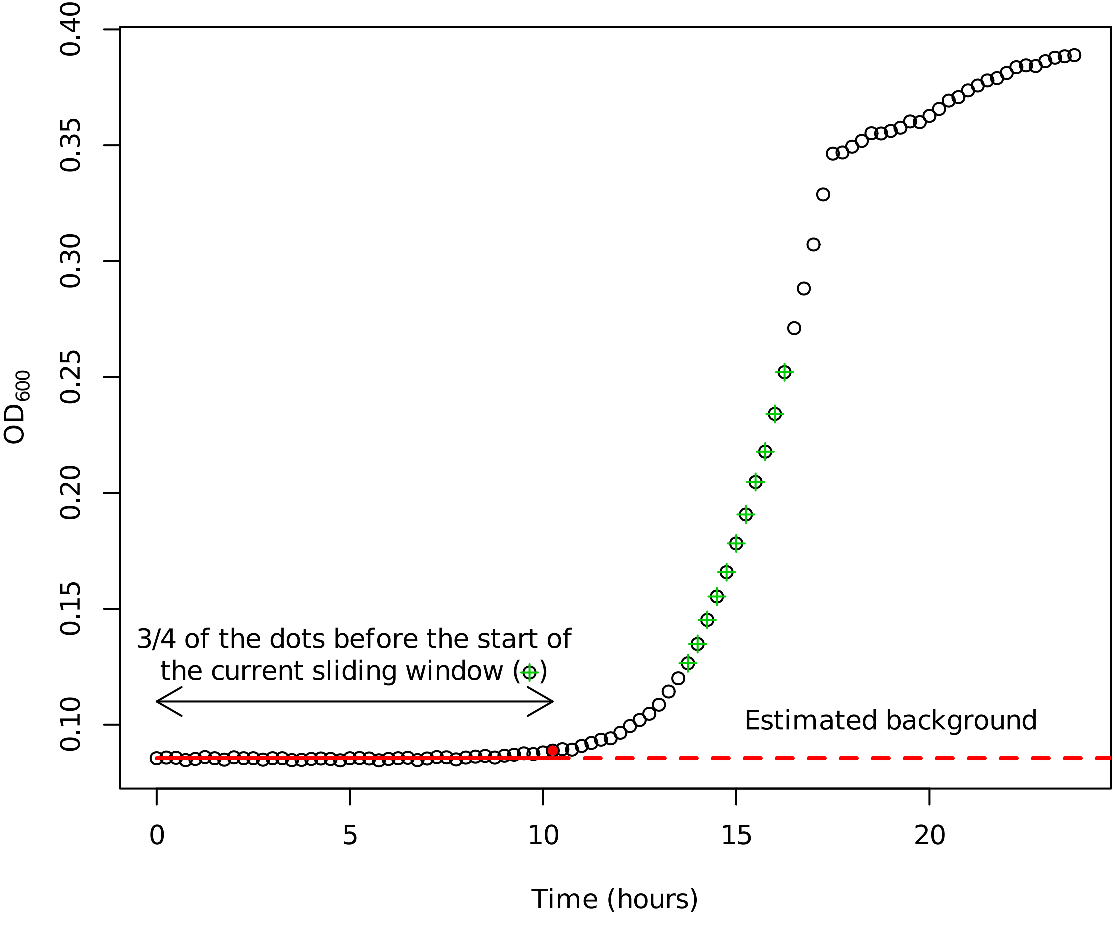

### heterogeneity between initial OD.pdf

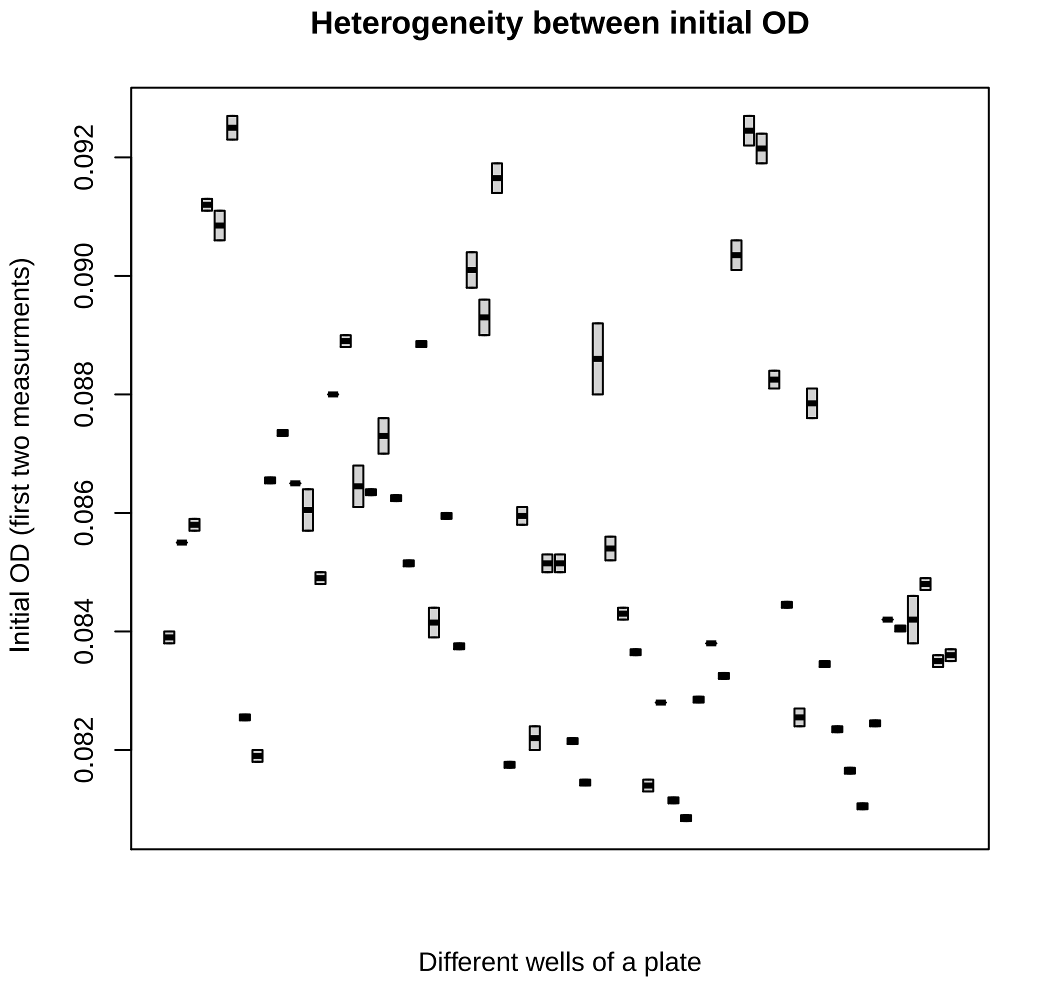
